# Fis1 is required for the development of the dendritic mitochondrial network in pyramidal cortical neurons

**DOI:** 10.1101/2025.01.07.631801

**Authors:** Klaudia Strucinska, Parker Kneis, Travis Pennington, Katarzyna Cizio, Patrycja Szybowska, Abigail Morgan, Joshua Weertman, Tommy L Lewis

## Abstract

Mitochondrial ATP production and calcium buffering are critical for metabolic regulation and neurotransmission making the formation and maintenance of the mitochondrial network a critical component of neuronal health. Cortical pyramidal neurons contain compartment-specific mitochondrial morphologies that result from distinct axonal and dendritic mitochondrial fission and fusion profiles. We previously showed that axonal mitochondria are maintained at a small size as a result of high axonal mitochondrial fission factor (Mff) activity. However, loss of Mff activity had little effect on cortical dendritic mitochondria, raising the question of how fission/fusion balance is controlled in the dendrites. Thus, we sought to investigate the role of another fission factor, fission 1 (Fis1), on mitochondrial morphology, dynamics and function in cortical neurons. We knocked down Fis1 in cortical neurons both in primary culture and *in vivo*, and unexpectedly found that Fis1 depletion decreased mitochondrial length in the dendrites, without affecting mitochondrial size in the axon. Further, loss of Fis1 activity resulted in both increased mitochondrial motility and dynamics in the dendrites. These results argue Fis1 exhibits dendrite selectivity and plays a more complex role in neuronal mitochondrial dynamics than previously reported. Functionally, Fis1 loss resulted in reduced mitochondrial membrane potential, increased sensitivity to complex III blockade, and decreased mitochondrial calcium uptake during neuronal activity. The altered mitochondrial network culminated in elevated resting calcium levels that increased dendritic branching but reduced spine density. We conclude that Fis1 regulates morphological and functional mitochondrial characteristics that influence dendritic tree arborization and connectivity.

## INTRODUCTION

Neurons are highly polarized cells that form distinct subcellular compartments termed dendrites, axons and somas. These domains allow neurons to compartmentalize chemical and electrical signals received from surrounding cells, integrate them, and pass them along in a circuit like manner. Neuronal polarization is established by cell-intrinsic and -extrinsic factors, including extracellular gradients, spatially-limited cytoskeletal components, and organelle localization and dynamics [1–4].

Mitochondria are one of the most abundant organelles in neurons, presumably a consequence of the neuron’s high energetic state and its necessity for precise calcium handling. Interestingly, mitochondrial morphology and abundance are strikingly different in the axons and dendrites of cortical pyramidal neurons. In the dendrites, mitochondria have an elongated tubular morphology with high occupancy of the dendritic space, while in the axon mitochondria are sparse and individual entities positioned at specific locations [5–7]. The neurons’ ability to maintain these compartmentalized mitochondrial morphologies appears to be critical for neuronal health as disruption due to environmental factors, genetic mutations or disease results in developmental disorders and potential for neurodegeneration [8–13].

Mitochondrial morphology is the result of the combined actions of mitochondrial biogenesis, trafficking, dynamics (fission & fusion), and removal (mitophagy) [14, 15]. Mitochondrial fission occurs following the recruitment of the GTPase Drp1 from the cytoplasm to the outer mitochondrial membrane by actin and ‘receptors’ localized at the surface of the mitochondrion [16–19]. In eukaryotic cells, four major Drp1 receptors have been identified: fission 1 (Fis1), mitochondrial fission factor (Mff), and mitochondrial elongation factors 1 and 2 (Mief1 & Mief2) [20–25]. While Drp1 dependent fission predominates, Drp1-independent fission has also been reported [26–28]. Fusion, which opposes fission, is carried out by the outer membrane localized GTPases Mfn1 and Mfn2, and the inner membrane localized GTPase Opa1 [10, 29].

Our recent work demonstrated that axonal mitochondria enter the axon and remain small as a consequence of a high rate of fission. This axonal fission relies mainly on mitochondrial fission factor (Mff)-mediated fission as loss of Mff activity results in elongated axonal mitochondria and altered axonal development [30]. Surprisingly, *Mff* knockdown had little effect on dendritic mitochondria morphology leaving several questions about the regulation of mitochondrial dynamics in the dendritic compartment of the neuron.

Fission 1 (Fis1) is the second most abundant Drp1 receptor in neurons [31, 32], but its role in eukaryotic mitochondrial fission and fusion remains controversial [20, 23, 33, 34]. Fis1p was the first fission factor identified in yeast, and along with Mdv4 makes up the only complex to recruit Drp1p [35, 36]. However, while knockout of *Fis1* in cell lines resulted in a small but significant increase in mitochondria size, when compared to *Drp1* or *Mff* knockout, Fis1 appears to play a minor role in eukaryotic mitochondrial morphology and may instead be more involved in mitophagy by end clipping [33, 37]. Interestingly, recent work shows that hFis1 can interface with both the fission and fusion machinery and may be important in adjusting the balance between mitochondrial fission and fusion [34]. While Fis1 is mainly known as a mitochondrial dynamics protein, it has also been proposed to play roles in apoptosis, mitochondrial motility, peroxisomal fission and regulating membrane contact sites [38].

In this manuscript, we explored the role of Fis1 in regulating compartment specific mitochondria morphology in cortical neurons. We report that reduced Fis1 activity surprisingly resulted in smaller dendritic mitochondria without affecting axonal mitochondria size. We show that mitochondrial motility is enhanced in Fis1 knockdown dendrites, resulting in an increased frequency of stochastic mitochondrial dynamics events, and thus lessening the fusion-dominant dendritic profile. Strikingly, the subcellular changes to the dendritic mitochondrial network perturbed cytosolic calcium handling that resulted in aberrant dendritic arborization and spine density.

## RESULTS

### Loss of Fis1 activity specifically affects the dendritic mitochondrial network

Previous work demonstrated that the Drp1 receptor mitochondrial fission factor (Mff) plays a critical role in the setup and maintenance of the axonal mitochondrial network [30]. While Mff is the most abundant Drp1 receptor in cortex and hippocampus according to publicly available RNA-seq data [31, 32], three other Drp1 receptors are present and their roles in the setup and maintenance of the neuronal mitochondrial network are mostly unexplored. To determine if Fis1, as the second most abundant receptor, has a role in compartmentalized neuronal mitochondria morphology, we first visualized mitochondrial Fis1 abundance on dendritic and axonal mitochondria in primary cultured mouse cortical neurons (**Supplementary Figure 1**). Staining for endogenous Fis1 revealed that Fis1 intensity is much higher on dendritic mitochondria when compared to axonal mitochondria even after accounting for mitochondrial size. This result suggested that Fis1 may play a more prominent role in mitochondrial dynamics of the dendrites than axons of cortical neurons.

To test the role of Fis1 in cortical neurons, we next validated commercially available shRNA mediated knockdown constructs for mouse *Fis1*. We identified two sequences (see Methods) that provided approximately 60 to 80% knockdown efficiency (**Supplementary Figure 2**). In order to visualize mitochondrial morphology and test the loss of Fis1 activity in cortical pyramidal neurons in a cell autonomous manner, we implemented either *ex utero* (EUE) or *in utero* (IUE) electroporation of plasmid DNA at E15.5 to sparsely manipulate progenitors giving rise to layer 2/3 cortical neurons [4, 30, 39, 40]. Following EUE or IUE of shRNA constructs for *Fis1*, along with plasmid DNA encoding a fluorescent cytoplasmic filler protein (tdTomato) and a mitochondria-targeted yellow fluorescent protein (mt-YFP) at E15.5 and visualization of neurons at 17DIV or P21, dendritic mitochondrial length was surprisingly decreased compared to control neurons both *in vitro* (4.3µm ± 0.2 for control, 2.4µm ± 0.1 for Fis1C, 2.3µm ± 0.2 for Fis1 386, **Figure 1a-b**) and *in vivo* (4.9µm ± 0.4 for control, 2.7µm ± 0.2 for Fis1C, 2.8µm ± 0.2 for Fis1 386, **Figure 1c-d**). This outcome is unlikely to be the result of an off-target effect as the two shRNAs have distinct targets, however to confirm the observed alteration in mitochondrial morphology is a result of Fis1 activity, we designed and subcloned a plasmid to express an shRNA impervious mouse Fis1 tagged with the hemagluttin (HA) tag (Fis1 imp, **Supplementary Figure 2**). After validating that it localized to mitochondria and was resistant to shRNA knockdown, we co-expressed a low level of the plasmid (0.33µg/µL) in combination with those above and were able to rescue the decrease in dendritic mitochondria length (3.9µm ± 0.1 for Fis1 rescue, (**Figure 1a**)). We also designed CRISPR guides to knockout mouse Fis1 (**Supplementary Figure 2**) and confirmed the loss of Fis1 resulted in reduced dendritic mitochondrial length ((3.4µm ± 0.1 for control, 2.4µm ± 0.2 for knockout (**Figure 1a-b**)). While Fis1 loss had a strong impact on dendritic mitochondrial size, loss of Fis1 activity did not alter axonal mitochondria size (1.22µm ± 0.05 for control, 1.26µm ± 0.06 for Fis1C, 1.21µm ± 0.06 for Fis1 386, 1.23µm ± 0.02 for CRISPR control, 1.23µm ± 0.03 for CRISPR KO, **Supplementary Figure 3**). These results show that Fis1 plays an important role in the formation of the compartmentalized mitochondrial network in neurons through the regulation of dendritic mitochondria morphology.

**Figure 1:**
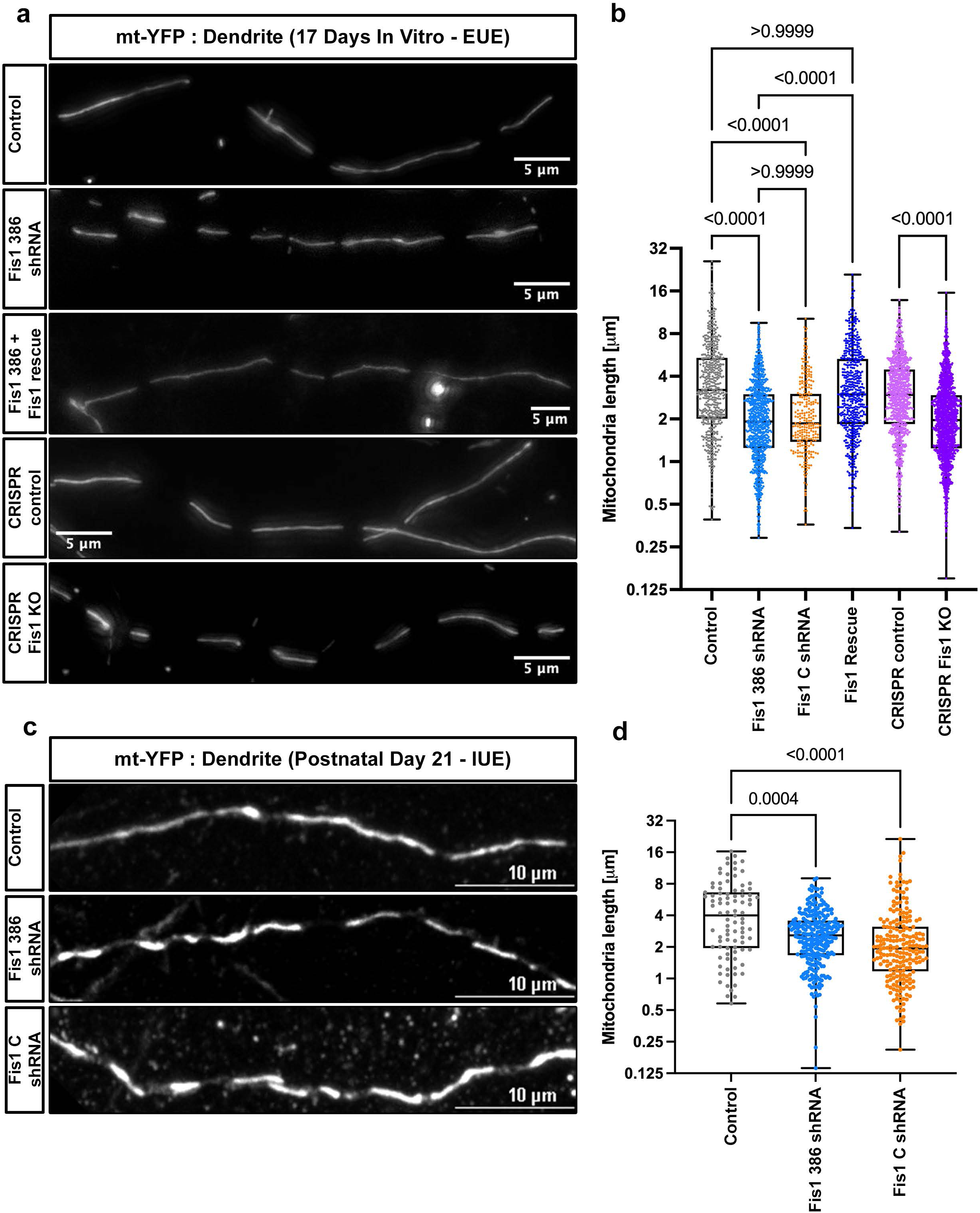
Fis1 knockdown results in shorter dendritic mitochondria both *in vitro* and *in vivo*. **a**) Representative dendritic segments from 17DIV cortical neurons showing mitochondrial morphology via matrix targeted YFP for each of the labeled conditions. **b**) Quantification of mitochondrial lengths for each of the indicated conditions demonstrating that loss of Fis1 results in shorter dendritic mitochondria. **c**) Representative dendritic segments from P21 layer 2/3 cortical neurons showing mitochondrial morphology via matrix targeted YFP for each of the labeled conditions in vivo. **d**) Quantification of mitochondrial lengths for each of the indicated conditions demonstrating that loss of Fis1 in vivo recapitulates the mitochondrial phenotype observed in cultured neurons. Control _in vitro_ = 546 mitochondria; Fis1 386 shRNA _in vitro_ = 873 mitochondria; Fis1 C shRNA _in vitro_ = 262 mitochondria; CRISPR control _in vitro_ = 949 mitochondria; CRISPR KO _in vitro_ = 1613 mitochondria; Control _in vivo_ = 91 mitochondria; Fis1 386 shRNA _in vivo_ = 272 mitochondria; Fis1 C shRNA _in vivo_ = 217 mitochondria. p values are indicated in the figure following Kruskal-Wallis tests. Data are shown as individual points on box plots with 25^th^, 50^th^ and 75^th^ percentiles indicated with whiskers indicating min and max values. Scale bars, 5 μm in a, 10 μm in c.

### Fis1 loss increases both dendritic mitochondria motility and dynamics

To characterize how Fis1 loss results in decreased mitochondria size, we employed the use of matrix-targeted, photo-activatable GFP [41] which allowed us to quantify the motility and frequency of mitochondrial fission and fusion in the dendrites of cortical neurons (**Figure 2**). Time-lapse imaging following photo-activation revealed that loss of Fis1 activity results in both dramatically increased mitochondrial motility and mitochondrial dynamics in the dendrites of cortical neurons. As neurons mature, mitochondria in both the axons and dendrites become fixed at specific points along the process leading to a large proportion of stationary mitochondria [42–45]. As expected in control dendrite segments at 14DIV, we observed that only ∼3 percent (2.5% ± 0.6) of mitochondria were motile over a 15 minute imaging session. However, upon Fis1 loss, the percent of motile dendritic mitochondria increased almost four times (11.6% ± 2.1 for Fis1 C, 13.3% ± 1.8 for Fis1 386, **Figure 2a-e**). The previously noted oscillating behavior of neuronal mitochondria [46] was also increased following Fis1 knockdown (**Figure 2d**). Concurrently, by quantifying the fission and fusion events of photoactivated mitochondria, we observed a doubling of mitochondrial fission (1.1 ± 0.1 events per 15 mins for control, 2.2 ± 0.2 for Fis1 C, 2.4 ± 0.2 for Fis1 386), as well as modestly increased fusion rates (1.6 ± 0.2 events per 15 mins for control, 1.9 ± 0.3 for Fis1 C, 2.9 ± 0.3 for Fis1 386) in the dendrites following the loss of Fis1 activity (**Figure 2f-g**) leading to an apparent loss of the fusion dominant phenotype observed in control neurons. These results demonstrate that Fis1 plays an important role in both dendritic mitochondrial motility and dynamics in cortical neurons.

**Figure 2:**
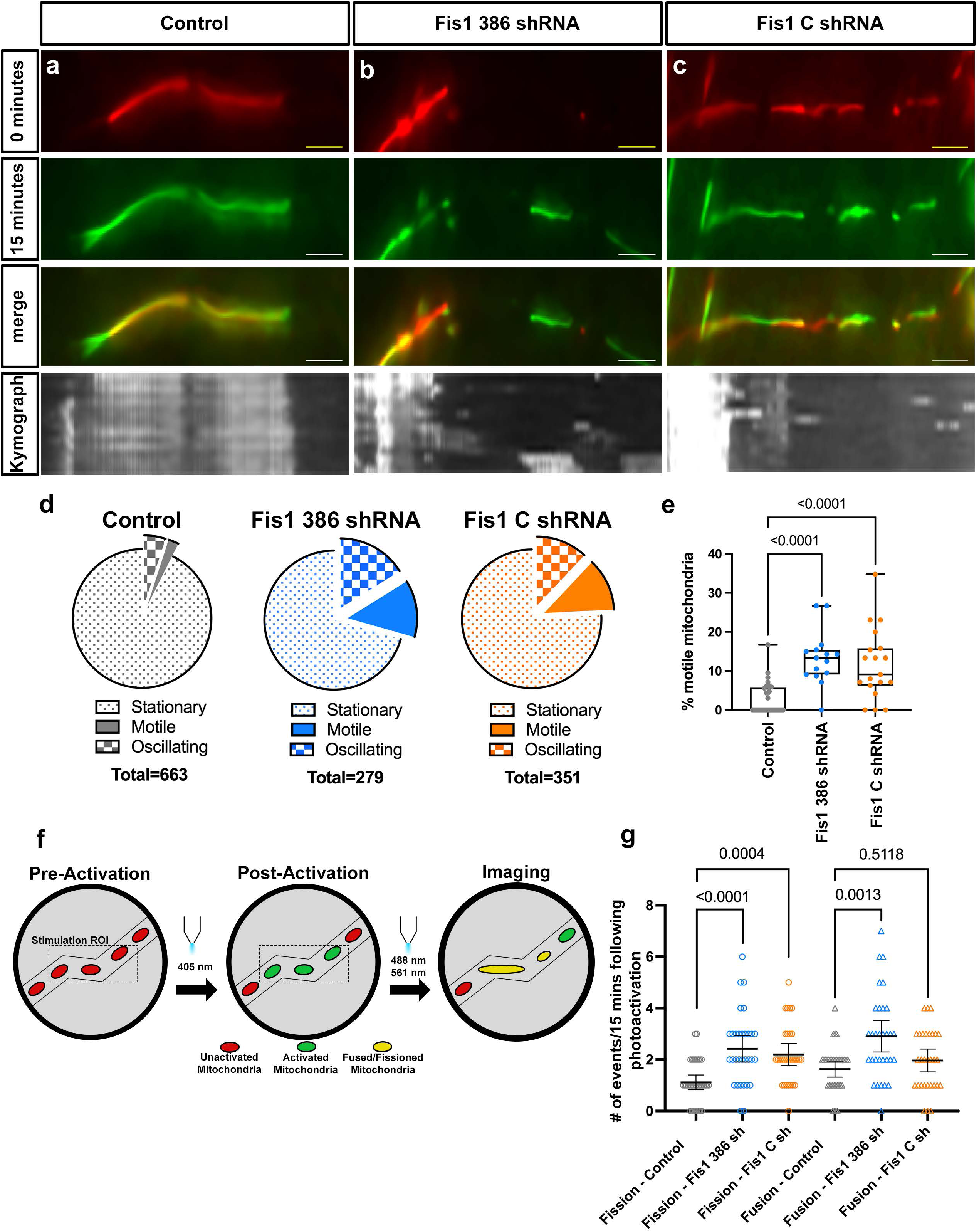
Loss of Fis1 activity increases both mitochondria motility and mitochondrial dynamics. **a**) Photo-activated mitochondria in a control dendrite at t=0 (red) and at t=15 minutes (green). The merged images showing overlap of the two timepoints and a kymograph of the entire 15 minute imaging session demonstrating little dendritic mitochondria movement in mature dendrites. **b,c**) Photo-activated mitochondria in a Fis1 386 shRNA (**b**) or Fis1 C shRNA (**c**) dendrite at t= 0 (red) and at t=15 minutes (green). The merge images showing overlap of the two timepoints and a kymograph of the entire 15 minute imaging session demonstrating increased mitochondria motility following Fis1 knockdown. **d**) Pie graphs showing percentages of stationary (dotted area), oscillating (checkered area), and motile (filled area) mitochondria in the dendrites of labeled conditions. **e**) Quantification of motile mitochondria percentage for dendrite segments demonstrating increased motility in Fis1 knockdown neurons. **f**) Scheme for photo-activation experiments to quantify fission and fusion dynamics in neuronal dendrites. **g**) Quantification of dendritic mitochondria fission and fusion rates as events per 15 minutes for control and knockdown dendrites showing that both fission and fusion are increased upon Fis1 loss in neurons. Control _motility_ = 37 segments, 663 mitochondria; Fis1 386 shRNA _motility_ = 15 segments, 279 mitochondria; Fis1 C shRNA _motility_ = 19 segments, 351 mitochondria; Control _dynamics_ = 35 segments, 39 fission events, 57 fusion events; Fis1 386 shRNA _dynamics_ = 31 segments, 75 fission events, 90 fusion events; Fis1 C shRNA _dynamics_ = 30 segments, 64 fission events, 57 fusion events. p values are indicated in the figure following Kruskal-Wallis tests. Data are shown as individual points on box plots with 25^th^, 50^th^ and 75^th^ percentiles indicated with whiskers indicating min and max values for e, or individual points with mean ± SEM for g. Scale bars, 5 μm.

### Fis1 knockdown impacts the functional parameters of dendritic mitochondria

To determine whether the altered morphology observed following Fis1 knockdown impacts the mitochondrion’s functional state, we performed EUE with DNA encoding either a control plasmid (pLKO) or the Fis1 386 shRNA and matrix targeted YFP at E15.5 and cultured to 14DIV. At 14DIV, we loaded the cultured neurons with tetramethylrhodamine methyl ester (TMRM) in order to visualize mitochondrial membrane potential. We observed slightly reduced uptake of TMRM into Fis1 knockdown dendritic mitochondria via signal intensity (680 ± 27 for control, 569 ± 34 for Fis1 386, **Figure 3a-c**) supporting a reduction in membrane potential. To confirm this observation, we performed EUE with DNA encoding the matrix targeted pH-sensitive fluorescent protein SypHer (mt-SypHer) and (mt-mScarlet) at E15.5. At 14 DIV, we measured the ratio of mt-SypHer to mt-mScarlet in dendritic mitochondria and observed a similar reduction in the ratio in Fis1 knockdown neurons compared to control (1.104 ± 0.01 for control, 1.025 ± 0.01 for Fis1 386, **Figure 3d**) representing the acidification of dendritic mitochondrial matrices in Fis1 knockdown. As matrix acidification is potentially associated with impaired proton pumping to the intermembrane space, we tested how dendritic mitochondria responded to complex 3 inhibition by antimycin A. Treatment with 1.25µM antimycin A led to a slightly larger reduction in TMRM signal in Fis1 knockdown mitochondria as compared to control dendritic mitochondria (−12.7% ± 2.0 for control, −23.0% ± 2.3 for Fis1 386, **Figure 3e-f**) demonstrating an increased sensitivity to complex 3 inhibition.

**Figure 3:**
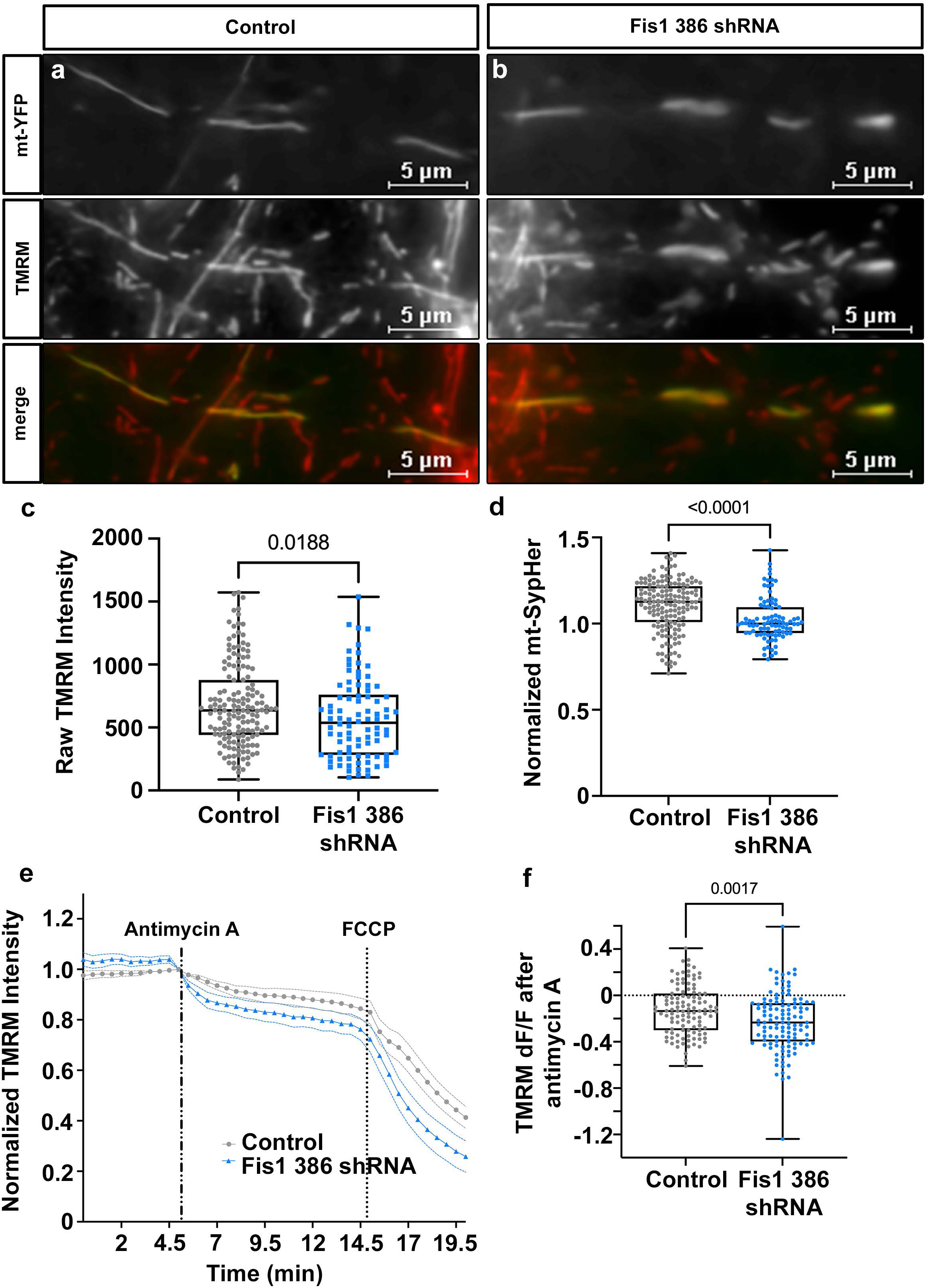
Dendritic Fis1 knockdown mitochondria have slightly reduced membrane potential and matrix pH. **a**) Representative images of dendritic mitochondria co-electroporated with mt-YFP (top) and control plasmid followed by loading with TMRM (middle). **b**) Representative images of dendritic mitochondria co-electroporated with mt-YFP (top) and Fis1 386 shRNA followed by loading with TMRM (middle). **c**) Quantification of raw TMRM dendritic mitochondria intensities showing reduced membrane potential in Fis1 knockdown mitochondria. **d**) Quantification of normalized mt-SypHer showing that Fis1 knockdown results in a slightly more acidic matrix pH. **e**) Graph of relative TMRM intensity following treatment with antimycin A and FCCP. **f**) Quantification showing dendritic mitochondria upon Fis1 loss are more sensitive to complex three inhibition. Control _TMRMinitial_ = 155 mitochondria; Fis1 386 shRNA _TMRMinitial_ = 90 mitochondria; Control _mtSypHer_ = 155 mitochondria; Fis1 386 shRNA _mtSypHer_ = 90 mitochondria; Control _TMRManta_ = 113 mitochondria; Fis1 386 shRNA _TMRManta_ = 117 mitochondria. p values are indicated in the figure following Mann-Whitney tests. Data are shown as individual points on box plots with 25^th^, 50^th^ and 75^th^ percentiles indicated with whiskers indicating min and max values for c,d & f, or mean ± SEM for e. Scale bars, 5 μm.

### Loss of Fis1 activity reduces dendritic mitochondria calcium uptake following evoked neuronal activity

Recent work has shown that dendritic mitochondria play an important role in calcium handling during neuronal activity [45, 47–50]. As alterations to mitochondrial membrane potential or matrix volume would impact mitochondrial calcium uptake, we tested if Fis1 knockdown had an impact on matrix calcium accumulation during glutamate uncaging which induces synaptically relevant calcium influx [51]. Following EUE at E15.5 with plasmids encoding mitochondrial matrix targeted GCaMP6f (mt-GCaMP6f) and mt-mScarlet, we imaged 14-16 DIV neurons following a single dendritic stimulation in imaging media supplemented with 1.3 mM MNI-glutamate. 2-3 seconds following stimulation, we observed a robust ∼3 fold increase in mt-GCaMP6f fluorescence for control neurons, while in Fis1 knockdown neurons the increase in fluorescence occurred on the same time scale but was reduced (2.7 fold ± 0.2 for control, 2.0 fold ± 0.1 for Fis1 386, **Figure 4a-d**), arguing that dendritic mitochondria have a reduced capability to take up calcium during neuronal activity following Fis1 loss.

**Figure 4:**
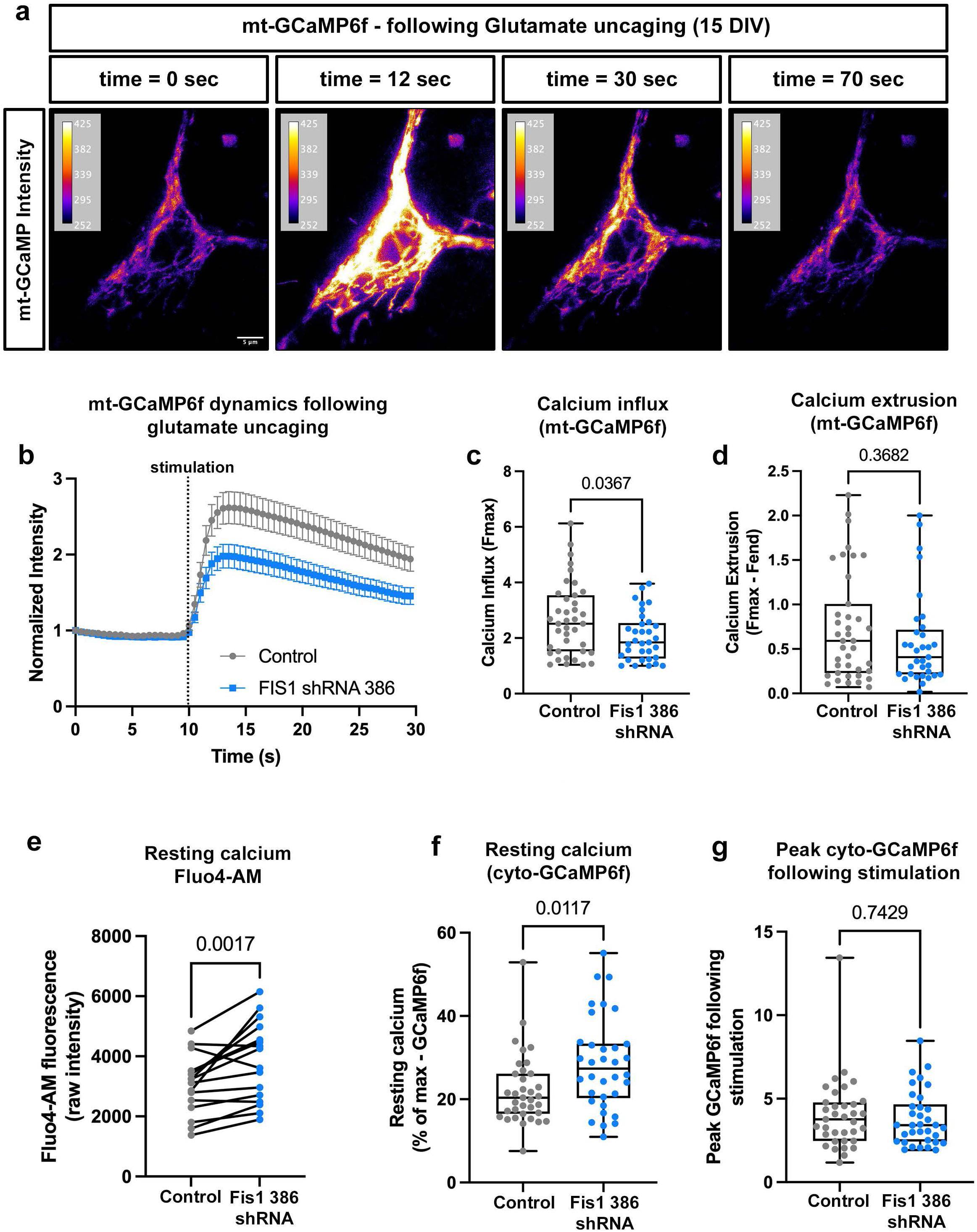
Fis1 loss reduces mitochondrial calcium uptake following evoked activity resulting in higher resting cytoplasmic calcium levels. **a**) Representative intensity images of somatodendritic mitochondria in a 15DIV neuron co-electroporated with mt-GCaMP6f (intensity gradient) and control plasmid during a time course following glutamate uncaging. **b**) Plot of normalized intensity of mt-GCaMP6f in control (grey) or Fis1 386 shRNA (blue) neurons following glutamate uncaging. **c**) F_max_ of mt-GCaMP6f following glutamate uncaging shows decreased calcium influx in Fis1 knockdown mitochondria. **d**) Quantification of mt-GCaMP6f extinction after F_max_ suggesting that Fis1 loss doesn’t affect mitochondrial calcium extrusion. **e**) Quantification of F_min_ Fluo4 showing increased resting cytoplasmic calcium levels in Fis1 knockdown neurons compared to surrounding wildtype neurons. **f**) Quantification of cytoplasmic GCaMP6f confirming increased resting calcium levels in Fis1 knockdown neurons. **g**) Peak GCaMP6f signals following glutamate uncaging show no difference in peak calcium levels in Fis1 knockdown neurons. Control _mtGCaMP6f_ = 39 dendrites; Fis1 386 shRNA _mtGCaMP6f_ = 33 dendrites; Control _Fluo4_ = 16 neurons; Fis1 386 shRNA _Fluo4_ = 16 neurons; Control _cytoGCaMP6f_ = 35 neurons; Fis1 386 shRNA _cytoGCaMP6f_ = 34 neurons. p values are indicated in the figure following Mann-Whitney tests (except e which is a Wilcoxon matched pairs test). Data are shown as individual points on box plots with 25^th^, 50^th^ and 75^th^ percentiles indicated with whiskers indicating min and max values, except for e which is a paired comparison of values for each pair. Scale bar, 5 μm.

### Decreased mitochondrial uptake of calcium increases resting cytoplasmic calcium

To understand the impact of reduced mitochondrial calcium uptake on neuronal calcium handling, we tested if cytoplasmic calcium levels are altered in Fis1 knockdown neurons. First, we loaded 14DIV cortical cultures with Fluo4-AM, a membrane permeable calcium sensing dye, to visualize resting calcium levels, and observed a significant increase in resting calcium levels in Fis1 knockdown neurons compared to nearby, non-electroporated neurons in the same dish (3053 au ± 247 for control, 3832 au ± 331 for Fis1 386, **Figure 4e**). Next, cytoplasmic GCaMP6f was used with the glutamate uncaging protocol to visualize calcium accumulation in the cytoplasm following evoked neuronal activity in cultured 14-16 DIV cortical neurons. Interestingly, the resting fluorescence levels of cytoplasmic GCaMP were higher in Fis1 knockdown neurons following normalization to the maximum fluorescence upon ionomycin addition (22.2% ± 1.4 for control, 28.6% ± 1.9 for Fis1 386, **Figure 4f**) confirming the results with Fluo4-AM, but peak levels of calcium influx were unaffected following stimulation (4.0 fold ± 0.4 for control, 3.8 fold ± 0.3 for Fis1 386, **Figure 4g**). These results argue that Fis1 loss does not impact peak calcium influx following a sufficient stimulus but that these neurons have a higher resting calcium which may result in a lower threshold for activation.

### Fis1 knockdown results in decreased ER calcium uptake following evoked activity

As mitochondrial transfer of calcium to endoplasmic reticulum (ER) is known to play an important role in calcium handling and calcium oscillations [47], we tested whether or not ER calcium dynamics were altered following Fis1 loss. Using the glutamate uncaging protocol on 14-16DIV cortical neurons electroporated with either control plasmid or Fis1 386 shRNA along with an ER localized and optimized GCaMP6 (erGCaMP6-150), we observed a trend towards slightly reduced ER calcium release (−0.11 ± 0.02 for control, −0.08 ± 0.01 for Fis1 386 shRNA), but significantly reduced ER calcium reuptake following stimulation in Fis1 knockdown neurons (0.12 ± 0.04 for control, 0.03 ± 0.02 for Fis1 386 shRNA, **Supplementary Figure 4**).

### Postsynaptic calcium responses are altered during spontaneous neuronal activity following Fis1 loss

Since Fis1 knockdown neurons have altered calcium handling and a higher resting calcium level, we imaged 14-16DIV cultured neurons electroporated with GCaMP6f and either control plasmid or Fis1 386 shRNA to determine if loss of Fis1 altered spontaneous neuronal activity. Following imaging at 2fps for 2 minutes, we didn’t observe any significant differences in the number of calcium transients or maximum spike amplitude (**Supplementary Figure 5a**). However, we did observe a shift towards slightly larger amplitudes overall arguing for a higher influx of calcium following presynaptic input (**Supplementary Figure 5b)**. We also observed an increase in the coefficient of variation in Fis1 knockdown neurons suggesting that the postsynaptic response to presynaptic input is less tightly controlled in these neurons (**Supplementary Figure 5c)**.

### Fis1 knockdown results in increased basal dendrite branching

As calcium regulation plays an important role in many cytoskeletal processes, and is well known to impact dendritic formation and branching, we set out to test whether loss of Fis1 activity results in altered dendritic branching [4, 52]. Thus, we performed Scholl analysis [53] on sparsely electroporated 14DIV cortical neurons electroporated with plasmids encoding tdTomato and either control plasmid or Fis1 386 shRNA. Interestingly, we found an increase in the number of crossings between 290 microns and 400 microns from the center of the cell body (**Figure 5a-c**) demonstrating an increase in the dendritic branching of Fis1 knockdown neurons.

**Figure 5:**
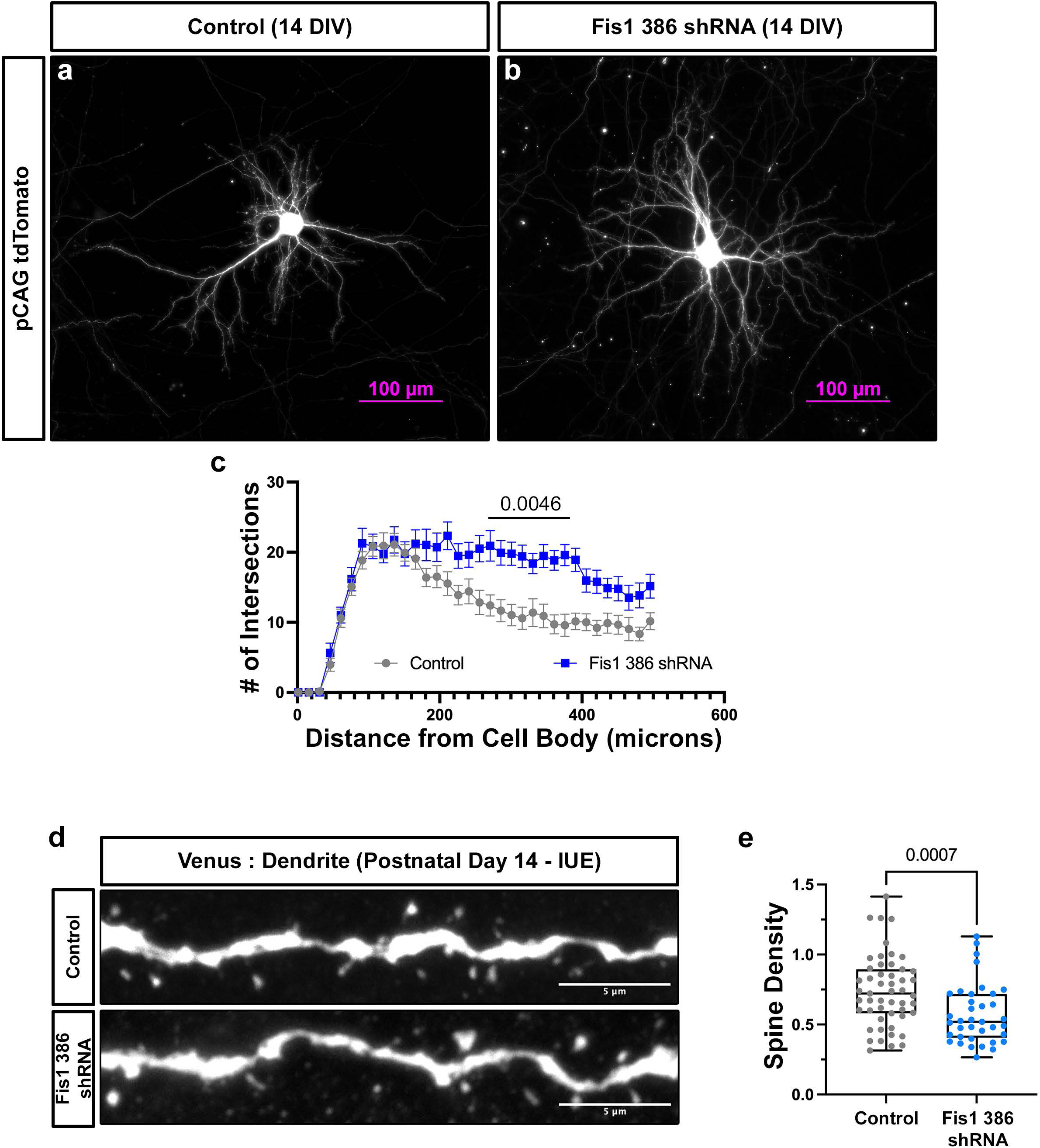
Dendritic branching and spine density are altered during development following Fis1 knockdown. **a-b**) Representative images of 14DIV cultured cortical neurons electroporated with tdTomato and either control (left, a) or Fis1 shRNA (right, b). **c**) Quantification of dendritic branching showing that Fis1 knockdown neurons have increased branching. **d**) Representative images of P14 layer 2/3 basal dendritic segments electroporated with Venus YFP and either control (top) or Fis1 shRNA (bottom) to visualize spine density. **e**) Quantification of dendritic spine density at P14 demonstrating that loss of Fis1 results in a reduced spine density *in vivo*. p values are indicated in the figure following Mixed-effects analysis (c), or Mann-Whitney test (e). Control _branching_ = 24 neurons; Fis1 386 shRNA _branching_ = 24 neurons. Control _spines_ = 51 segments; Fis1 386 shRNA _branching_ = 37 segments. Data are shown as a plot of the number of crossings per distance from the cell body (b), or individual points on box plots with 25^th^, 50^th^ and 75^th^ percentiles indicated with whiskers indicating min and max values (d). Scale bars, 100 μm for a, 5 μm for c.

### Dendritic spine density is decreased following Fis1 knockdown *in vivo*

As dendritic branching was increased, we tested if Fis1 knockdown also affected dendritic spine density. To maximize visualization of individual spines, we sparsely labeled neurons by IUE at E15.5 with a Flpe-dependent GFP plasmid and either control plasmid or Fis1 386 shRNA plasmid. At P14, we perfused the mice, sectioned and stained the brains, then imaged basal dendrites of layer 2/3 cortical pyramidal neurons at high magnification. Upon knockdown of Fis1, we observed a significant reduction in the density of spines on basal dendrites during development *in vivo* (0.75 spines/µm ± 0.04 for control, 0.57 spines/µm ± 0.03 for Fis1 386 shRNA, **Figure 5d-e**). This result coupled with the branching results above demonstrates that loss of Fis1 results in altered dendritic development.

## DISCUSSION

In the present study, we reveal that Fis1 activity is critical in the formation and maintenance of the dendritic mitochondrial network. Our results demonstrate that loss of Fis1 surprisingly reduces dendritic mitochondrial length by increasing both the rate of trafficking and dynamics along the dendrites thus tipping the balance away from a fusion dominant phenotype in the dendrites. The resulting mitochondria have lower membrane potential, increased sensitivity to antimycin A, and a reduced capacity to take up calcium during neuronal activity. Fis1 loss also resulted in increased dendritic branching and reduced spine density presenting a scenario whereby reduced calcium handling by the mitochondrial network alters dendritic development.

Our results argue for a compartment-specific role for Fis1 in the development of the dendritic mitochondrial network as the loss of Fis1 activity alters dendritic mitochondrial size, trafficking and dynamics without altering the axonal mitochondrial pool. The question of how Fis1 specifically regulates dendritic mitochondria remains unclear. Our immunocytochemistry data indicates that Fis1 protein is nearly twice as abundant on dendritic mitochondria than axonal mitochondria likely explaining the preferential impact of its loss on the dendritic pool of mitochondria. Surprisingly though, the loss of Fis1 resulted in decreased, not increased, mitochondrial length. This could potentially arise from a few different possibilities: (1) Fis1 may interact with other ‘Drp1 receptors’ such as Mff or Mief1/2 to alter the recruitment or activity of Drp1 at the outer mitochondrial membrane [23–25, 54], (2) Fis1 may preferentially recruit non-active conformations of Drp1 to dendritic mitochondria [14, 19, 37, 55], or (3) human Fis1 was recently shown to interact with the fusion machinery thus the loss of Fis1 may directly impact fusion processes [34]. Each of these scenarios could support a model where higher Fis1 levels in the dendrites results in increased mitochondrial length, but future work will be necessary to test these possibilities.

Strikingly, Fis1 loss resulted in both increased mitochondrial trafficking and increased mitochondrial dynamics which may suggest a mechanism where Fis1 is a component of the mitochondrial anchoring system. Previous work from multiple groups has shown that mitochondrial trafficking is reduced with dendritic maturation and mitochondria become stably captured at specific points along the dendrites [44, 45, 50] but the mechanism has remained elusive. Coupled with our results that Fis1 impacts mitochondrial-ER calcium coupling, it is plausible that Fis1 may have a potential role in MERCS formation or dynamics leading to a model where mitochondrial capture and the development of mitochondrial-ER coupling in the dendrites is linked. In fact, recent work in non-neuronal cells have shown that Fis1 is present at MERCS [19, 56, 57] but its exact role there remains unclear. If Fis1 plays a role in mitochondrial capture, it will be interesting to determine if the mere reduction in mitochondrial motility is sufficient to dampen mitochondrial dynamics in a manner that favors a pro-fusion phenotype.

The role of calcium as a cytosolic signal for dendritic branching and postsynaptic development is well established [4, 52]. Even before evoked activity is present, developing neurons have spontaneous calcium transients [58] that coincide with the timing of dendritic branching. A major pathway connecting dendritic development to calcium signaling is via calmodulin and calcium/calmodulin-dependent protein kinases (CaMKs) [59]. CaMKII is the most well studied with CaMKII alpha and beta shown to alter dendritic outgrowth, filopodial extension, and regulate interaction with actin [60–62]. Dendritic spine growth during long-term potentiation (LTP) is also linked to calcium dynamics through the remodeling of spine actin [63–65]. Interestingly, a number of groups have recently observed that synaptic activity is directly tied to mitochondrial fission via calcium signaling to directly link neuronal activity to metabolic and calcium handling needs [49, 66–68]. Future investigation will be necessary to determine if Fis1 plays a role, either directly or indirectly with Mff, in this dendritic calcium-dependent fission.

Taken together, our data demonstrate that Fis1 plays a compartment-specific, yet unconventional, role in the development of the dendritic mitochondrial network.

## Author Contributions

Conceptualization: TL. Formal analysis: KS, TP, PK, AM, TL. Funding acquisition: TL. Investigation: KS, TP, PK, PS, JW, KC. Methodology: KS, TP, PK, TL. Visualization: KS, TP, PK, TL. Writing – original draft: KS, TL. Writing – review & editing: KS, TP, PK, TL.

## Acknowledgements

We thank past and present members of the Lewis lab, Julien Courchet, Yusuke Hirabayashi, and Seok-Kyu Kwon for feedback and discussion over the course of the project. We thank the OMRF Imaging core for excellent imaging support, and the OMRF vivarium staff excellent animal care. This research was supported by grants from NIGMS (R35GM137921) and the Presbyterian Health Foundation to TL.

## METHODS

### Animals

Animals were handled according to Institutional Animal Care and Use Committee (IACUC) approved protocols at the Oklahoma Medical Research Foundation (OMRF, 20-23, 23-18). Time-pregnant females of CD-1 IGS strain (Strain Code: 022) were purchased at Charles River Laboratories and used for *in utero* electroporation experiments and primary neuronal cultures.

### Plasmids

pCAG:mtYFP-P2A-tdTomato, pCAG:tdTomato and pCAG:mt-YFP were previously published in [69]. pCAG:2xmtpaGFP p2a 2xmtmScarlet was created by cloning a gene block encoding (from IDT) 2xmtpaGFP p2a 2xmtmScarlet into pCAG via restriction digest. pCAG:mt-SypHer was created by PCR of mt-SypHer from Addgene plasmid 48251 (a gift from Nicolas Demaurex) and cloning it 3’ to the CAG promoter. pCAG GCaMP6f was created by PCR of GCaMP6f from Addgene plasmid 40755 (a gift from Douglas Kim) and cloning it 3’ to the CAG promoter. pCAG mt-GCaMP6f was created by excising the YFP from pCAG mt-YFP and inserting GCaMP6f in its place. pCAG erGCaMP6-150 was created by PCR of GCaMP6-150 from Addgene plasmid 91777 (a gift from Douglas Kim) and cloning it 3’ to the CAG promoter while adding a KDEL retention motif to the C-terminus via the primers. pCAG HA-mouse Fis1 and pCAG HA-mouse Fis1 impervious were created by ordering gene blocks encoding a HA tag N-terminal to mouse Fis1 variant 1 or a mouse Fis1 variant 1 with the following sequences mutated (site 1 for Fis1 386 shRNA: cctggttcgaagcaaatacaa to tgccttgtgaggtctaagtacaa and site 2 for Fis1 C shRNA: ccaaggagctggaacgcctgattgataag to caaaggaacttgagcggctcatagataaa) and cloning them into the pCAG vector via restriction digest. pLKO.1 cloning vector was purchased from Addgene (plasmid 10878, a gift from David Root) and used as a control plasmid. Fis1 386 shRNA plasmid was ordered from Sigma (pLKO.1 TRCN0000124386, target sequence: CCTGGTTCGAAGCAAATACAA). Fis1 C shRNA plasmid was ordered from Origene (pGFP-C-shLenti clone C, target sequence: CCAAGGAGCTGGAACGCCTGATTGATAAG). Fis1 CRISPR guides were generated using CHOPCHOP online software targeting exon 3, 4 or 5. Oligos of each selected guide were ordered through IDT and cloned into pOrange (Addgene plasmid 131471) between Bbs1. Exon 3 guide: CGTGGGCAACTACCGGCTCAAGG, Exon 4 guide: AAGGCTCTAAAGTATGTGCGAGG, Exon 5 guide: CTGGTAGGCATGGCCATCGTTGG. CRISPR guide efficiency was tested using pCAG-EGxxFP (Addgene plasmid 50716) with the genomic DNA for the 3^rd^ thru 4^th^ exon of mouse Fis1 cloned between Nhe1 and EcoR1.

### Cell Lines

Human embryonic kidney cells (HEK293T/17) were purchased from ATCC (CRL-11268). 1×10^5^ cells were suspended in media (DMEM, Gibco) with penicillin/streptomycin (0.5x; Gibco) and FBS (Sigma) were seeded in 6 well tissue culture dishes (Falcon). Transfection with plasmid DNA (1 mg/mL) using jetPRIME® reagent (Polyplus) according to manufacturer protocol was performed 24 hours after seeding. For the EGxxFP assay cells were live imaged 48 hours after transfection. For western blotting 72 hours following transfection, cells were carefully washed with 1xPBS (Gibco) then collected into RIPA buffer with protease inhibitor cocktail.

### Western Blotting

Aliquots of the collected samples were separated by SDS-PAGE and then transferred to a polyvinylidene difluoride (PVDF) membrane (Amersham). After transfer, the membrane was washed 3X in Tris Buffer Saline (10 mM Tris-HCl pH 7.4, 150 mM NaCl) with 0.1% of Tween 20 (T-TBS), blocked for 1 hr at room temperature in Odyssey Blocking Buffer (TBS, LI-COR), followed by 4°C overnight incubation with the appropriate primary antibody in the above buffer. The following day, the membrane was washed 3X in T-TBS, incubated at room temperature for 1 hr with IRDye secondary antibodies (LI-COR) at 1:10,000 dilution in Odyssey Blocking Buffer (TBS), followed by 3X T-TBS washes. Visualization was performed by quantitative fluorescence using an Odyssey CLx imager (LI-COR). Signal intensity was quantified using Image Studio software (LI-COR). Primary antibody used for Western-blotting was Rabbit anti-HA (CST3724) or Rabbit anti-GAPDH (PA1-16777). Total protein was assessed using the Revert 700 Total protein stain (LI-COR 926-11010).

### *Ex utero* electroporation

A mix of endotoxin-free plasmid preparation (1-2 mg/mL) and 0.5% Fast Green (Sigma) mixture was injected using FemtoJet 4i (Eppendorf) into the lateral ventricles of isolated heads of E15.5 mouse embryos. Embryonic neural progenitor cells were electroporated using an electroporator (ECM 830, BTX) and gold paddles with four pulses of 20 V for 50 ms with 500 ms interval and an electrode gap of 1.0 mm. Dissociated primary neuron culture was performed after *ex utero* electroporation.

### Primary neuronal culture

Following *ex utero* electroporation, embryonic mouse cortices (E15.5) were dissected in Hank’s Buffered Salt Solution (HBSS) supplemented with Hepes (2.5 mM), CaCl_2_ (1 mM, Sigma), MgSO_4_ (1 mM, Sigma), NaHCO_3_ (4mM, Sigma) and D-glucose (30 mM, Sigma), hereafter referred to as cHBSS, and incubated in cHBSS containing papain (Worthington; 14 U/mL) and DNase I (100 μg/mL) for 15 min at 37 °C with a gentle flick between incubation. Samples were washed with cHBSS three times, and dissociated by pipetting on the fourth wash. Cells were counted using Countess™ (Invitrogen) and cell suspension was plated on poly-D-lysine (1 mg/mL, Sigma)-coated glass bottom dishes (MatTek) in Neurobasal media (Gibco) containing FBS (2.5%) (Sigma), B27 (1 X) (Gibco), and Glutamax (1 X) (Gibco). After 7 days, media was changed with supplemented Neurobasal media without FBS.

### *In utero* electroporation

A mix of endotoxin-free plasmid preparation (1-2 mg/mL) and 0.5% Fast Green (Sigma) was injected into one lateral hemisphere of E15.5 embryos using FemtoJet 4i (Eppendorf). Embryonic neural progenitor cells were labelled using the electroporator (ECM 830, BTX) with gold paddles at E15.5. Electroporation was performed by placing the anode (positively charged electrode) on the side of DNA injection and the cathode on the other side of the head. Five pulses of 38 V for 50 ms with 500 ms interval and an electrode gap of 1.0mm were used for electroporation. Embryos were randomly assigned to control or knockdown without regard to sex.

### Intracardial perfusion

Animals were anesthetized using 5% isoflurane mixed with air and exsanguinated 14 or 21 days after birth (P14 or P21) by terminal intracardial perfusion. Fixative was kept on ice during the entire procedure. Direct perfusion was performed with 30 mL of fixative: 2% PFA/0.075% GA in PBS. Animals were then dissected to isolate brains, that were later subjected to 20h post fixation in the same fixative that was used for perfusion.

### Immunocytochemistry

Cultured neurons were fixed for 10 min on ice in 2% PFA (Alfa Aesar) with 0.075% GA (Electron Microscopy Science, EMS), and then washed with PBS (Sigma). Cells were permeabilized with 0.2% Triton X-100 in PBS and incubation in 0.1% BSA and 2.5% goat serum in PBS was followed to block nonspecific signals. Primary and secondary antibodies were diluted in the blocking buffer described above and incubated at 4°C overnight. Coverslips were mounted on slides with Fluoromount G (SouthernBiotech). Primary antibodies used in these experiments were mouse anti-HA (Biolegend, 1:500), chicken anti-GFP (Aves Lab 1:1000), rabbit anti-dsRed (Abcam 1:1000), rabbit anti-Fis1 (Proteintech 1:100) and all secondary antibodies were Alexa-conjugated (Invitrogen) and used at 1:1000 dilution. For visualizing endogenous Fis1, after fixation the cultures were washed with PBS with 22mg/mL of glycine (Sigma) for 10 minutes. Cells were then placed in blocking buffer for 1 hour with 0.2% Saponin (Sigma) in 1xPBS with 5% NGS and 22mg/mL glycine. Primary (1:100 O/N at 4C) and secondary (1:500 2hrs at RT) antibodies were incubated in 1xPBS with 1% BSA (Sigma), and 0.2% Saponin.

### Immunohistochemistry

Following fixation and washing, brains were embedded in 3% low melt agarose (RPI, A20070) in 1x PBS. Brains in agarose cubes were sectioned using a vibratome (Leica VT1200) at 120 μm. Sections were then incubated with primary antibodies (chicken anti-GFP Aves Lab 1:1000, rabbit anti-dsRed Abcam 1:1000) that were diluted in the Blocking buffer (1%BSA, 0.2%TritonX-100, 5%NGS in PBS) at 4 °C for 48h. Subsequently sections were washed 6 times for 10 min in PBS and incubated with secondary antibodies (Alexa conjugated goat anti-chicken488 and goat anti-rabbit568 1:1000) at 4 °C for 48h. The excess of secondary antibodies was removed by six, 10 minutes washes in 1x PBS. Sections were then mounted on slides and coverslipped with Aqua PolyMount (PolyMount Sicences, Inc.) and kept at 4 °C.

### Live imaging of cultured neurons

Cultured neurons were imaged on a Nikon Ti2 widefield system equipped with a Hammamatsu ORCA-Fusion CMOS camera, a custom penta-band cube for 378/474/554/635/735 excitation with an Aura III light engine, and 60x (1.4NA) oil objective, and live imaging chamber from OXO (UNO-T-H-CO2) with objective warmer. In addition, a 405nm laser (LUN-F, 50mW) with XY galvo control (Opti-microscan) is connected to perform targeted ROI based stimulation. The whole system is controlled by Nikon Elements. For live imaging, culture medium was removed and replaced with warmed cHBSS. For each experimental paradigm both controls and experimental conditions were imaged with the same light powers, camera exposures and all other microscope settings.

For photo-activation experiments, 8 micron ROIs were selected along multiple dendrites of each cell imaged. A before stimulation image was taken, then each box was individually stimulated at 2% laser power for 100µs, immediately followed by timelapse imaging every 30 seconds for 15 minutes.

For TMRM experiments, neuron cultures were preloaded with 20nM TMRM for 20 minutes then placed in cHBSS with 5nM TMRM. Cells were allowed to equilibrate for 20 additional minutes before imaging. Antimycin A and FCCP (Sigma) were used at 1.25µM and added at the timepoints indicated while timelapse imaging every 30 seconds.

For Fluo4-AM experiments, neuron culture were loaded with 5µM Fluo4-AM (Life Technologies Corp, F14217) in imaging media for 20 minutes then imaged live.

For glutamate uncaging experiments, imaging media was supplemented with 1.3mM MNI-glutamate (MNI-caged-L-glutamate, Tocris 1490) and 0.001mM TTX. After an appropriate dendrite was identified and ROI masked, timelapse imaging was performed at 2fps for 30 seconds. At 10 seconds, the 405 laser (2% laser power for 100µs) was activated to stimulate the selected ROI.

For spontaneous activity, neuronal cultures in cHBSS were imaged at 2fps for 30 seconds to visualize GCaMP activity.

### Imaging of fixed brain sections and cultured neurons

Fixed samples were imaged on a Zeiss LSM 880 confocal microscope controlled by Zeiss Black software. Imaging required two lasers 488nm and 561nm together with Zeiss objectives 40x (1.2NA) with 2x digital zoom, or 100x oil (1.25NA) with 3x digital zoom.

### Quantification and statistical analysis

For data in figure 1 and supplementary figure 3, individual mitochondria were measured via the mt-YFP signal from isolated, secondary dendrite segments or axonal segments with NIS Elements AR (Nikon) using the length measurement tool on the raw images.

For data in figure 2, photo-activated dendritic mitochondria were manually tracked in NIS Elements AR to calculate motility and fission/fusion dynamics. Motility was assessed as movement greater than 5 microns, oscillating mitochondria are those that moved but less than 5 microns, while stationary mitochondria remained in place throughout the timelapse. Fission was counted if a single mitochondria split into two or more individual mitochondria, while fusion was counted by the mixing of matrix content resulting in an altered green to red ratio.

For data in figure 3, regions of interest were drawn around individual mitochondria in labeled dendritic segments. Average ROI intensity was collected in NIS Elements AR (Nikon).

For data in figures 4 and supplementary figures 4&5, regions of interest were drawn over the indicated areas of neurons. Average ROI intensity was collected over the time segments indicated via the Time Measurement plugin in NIS Elements AR. Calculations used to visualize/normalize the data (such as **Δ**F/F_0_) are presented on the y-axis of each graph in the figures.

For data in figure 5, branching was measuring by using the Scholl analysis plugin in FIJI on optically isolated 14DIV cortical neurons. Crossings were plotted and analyzed in GraphPad Prism. For spine data, 2 random basal dendrites per P14 neuron was imaged at high magnification. In NIS Nikon Elements AR, individual spines were manually counted and the length of the dendritic segment was calculated. Spine density was then quantified as spines per micron of dendrite.

Statistical analysis was done in GraphPad’s Prism 9. Statistical tests, p-values, and (n) numbers are presented in the figure legends. Gaussian distribution was tested using D’Agostino & Pearson’s omnibus normality test. All analyses were performed on raw imaging data without any adjustments. Images in figures/movies have been adjusted for brightness and contrast (identical for control and experimental conditions in groups compared), and have been processed with Nikon’s proprietary denoise.ai for visualization purposes only.

---

**Supplementary Figure 1:**
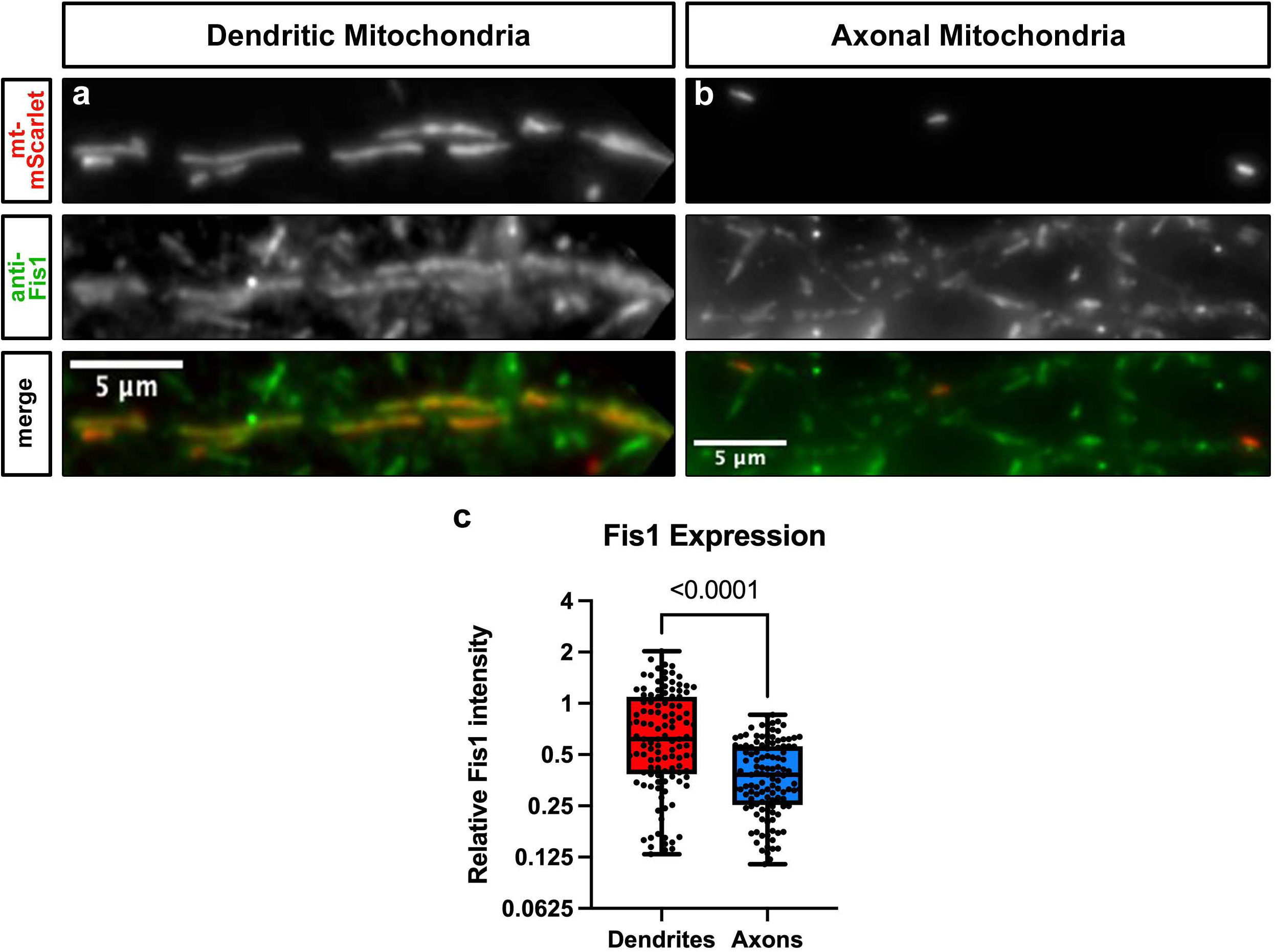
Fis1 protein levels are higher on dendritic mitochondria than axonal mitochondria. **a**) Representative images of dendritic mitochondria (red), and their overlap with endogenous Fis1 (green). **b**) Representative images of axonal mitochondria (red), and their overlap with endogenous Fis1 (green). **c**) Quantification of relative Fis1 intensity levels showing that dendritic mitochondria have more Fis1 associated with them than do axonal mitochondria. p value is indicated in the figure following a Mann-Whitney test. Dendrites = 115 mitochondria; Axons = 117 mitochondria. Data are shown as individual points on box plots with 25^th^, 50^th^ and 75^th^ percentiles indicated with whiskers indicating min and max values. Scale bars, 5 μm.

**Supplementary Figure 2:**
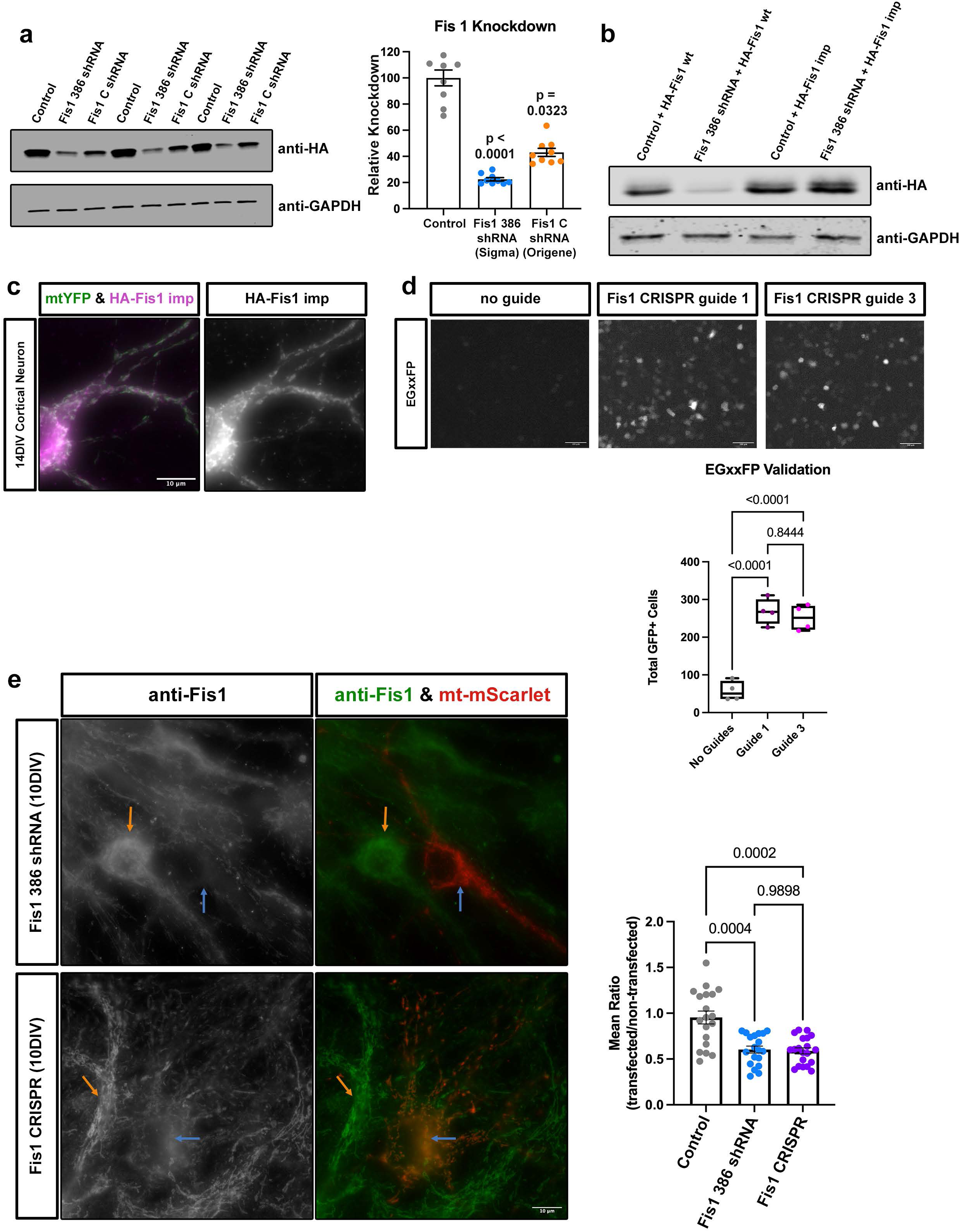
Fis1 shRNA and CRISPR validation. **a**) Representative images of western blots for knockdown of HA-tagged Fis1 by the indicated shRNA constructs with Gapdh as a loading control to demonstrate that both shRNA constructs effectively knockdown Fis1. **b**) Representative images of western blots to demonstrate that the HA-Fis1 impervious construct is not knocked down by the shRNA. **c**) Representative images of a 14DIV cortical neuron electroporated with mt-YFP (green) and HA-Fis1 impervious (magenta) to show that HA-Fis1 impervious expresses and is largely co-localized with mitochondria. **d**) Representative images of EGxxFP expression following cutting with Fis1 CRISPR guides demonstrating effective cutting. **e**) Representative images showing that both the shRNA and CRISPR constructs result in a significant reduction in endogenous Fis1 following electroporation compared to un-electroporated neighboring cells. p value is indicated in the figure following a Kruskal-Wallis test in a, or Brown-Forsythe and Welch ANOVA test in d,e. Control _in a_ = 9 wells; Fis1 386 shRNA _in a_ = 9 wells; Fis1 C shRNA _in a_ = 9 wells. No guide _in d_ = 4 wells; Fis1 guide exon 3 _in d_ = 4 wells; Fis1 guide exon 4 _in d_ = 4 wells. Control _in d_ = 19 cells; Fis1 386 shRNA _in d_ = 19 cells; CRISPR KO _in d_ = 20 cells. Data are shown as scatter plots with bar showing mean with SEM. Scale bars, 10 μm in c,e; 100 μm in d.

**Supplementary Figure 3:**
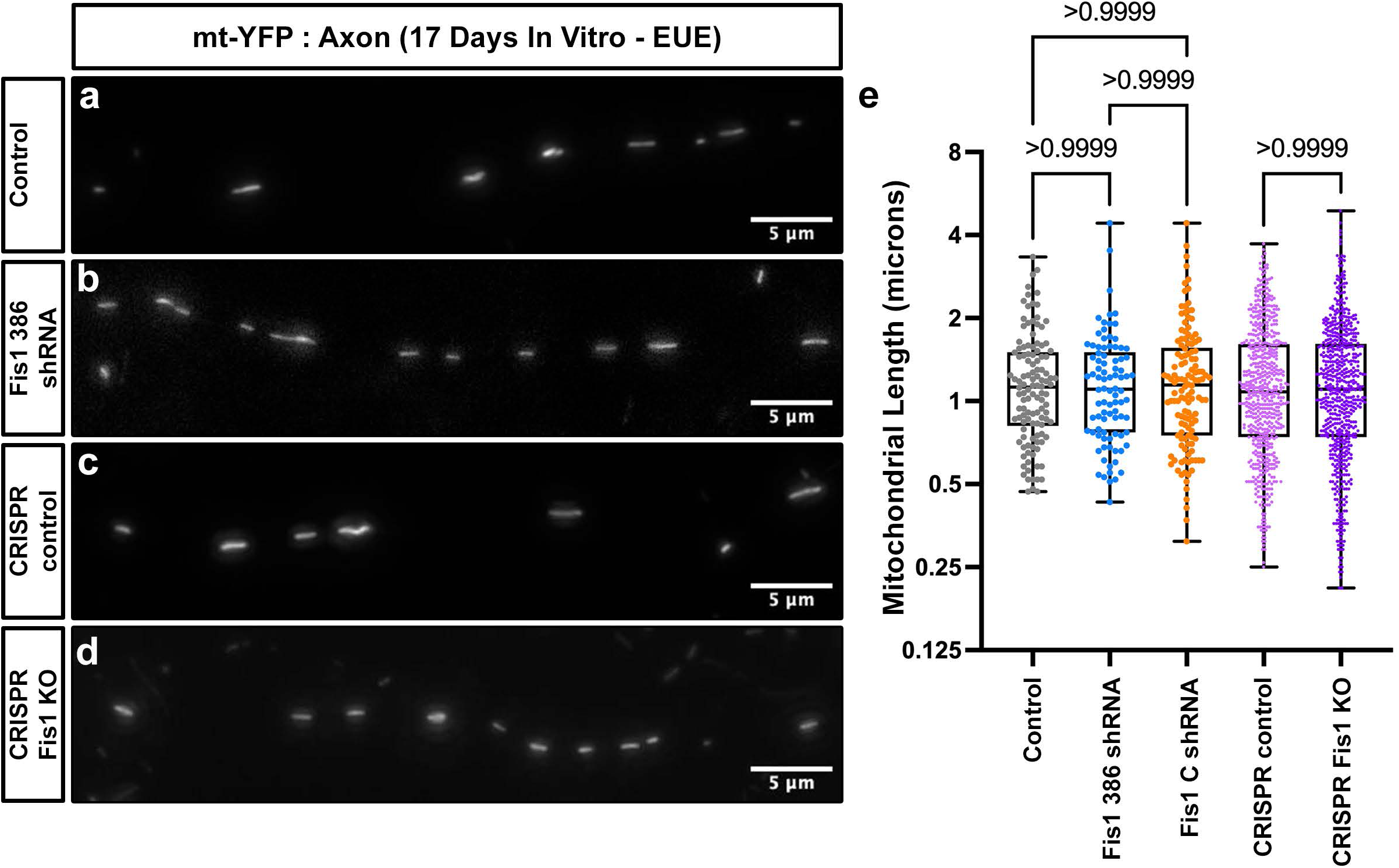
Fis1 knockdown doesn’t affect axonal mitochondria size. **a**) Representative images of axonal mitochondria in a 17DIV neuron co-electroporated with mt-YFP and control plasmid. **b**) Representative images of axonal mitochondria in a 17DIV neuron co-electroporated with mt-YFP and Fis1 386 shRNA. **c**) Representative images of axonal mitochondria in a 17DIV neuron co-electroporated with mt-YFP and CRISPR control plasmid. **d**) Representative images of axonal mitochondria in a 17DIV neuron co-electroporated with mt-YFP and CRISPR Fis1 KO plasmids. **e**) Quantification of mitochondrial length showing that neither Fis 1 knockdown construct alters axonal mitochondrial length. Control _axonal mitochondria_ = 22 segments, 118 mitochondria; Fis1 386 shRNA _axonal mitochondria_ = 14 segments, 90 mitochondria; Fis1 C shRNA _axonal mitochondria_ = 21 segments, 130 mitochondria; Control CRISPR _axonal mitochondria_ = 53 segments, 621 mitochondria; CRISPR Fis1 KO _axonal mitochondria_ = 54 segments, 638 mitochondria. p values are indicated in the figure following Kruskal-Wallis tests. Data are shown as individual points on box plots with 25^th^, 50^th^ and 75^th^ percentiles indicated with whiskers indicating min and max values. Scale bars, 5 μm.

**Supplementary Figure 4:**
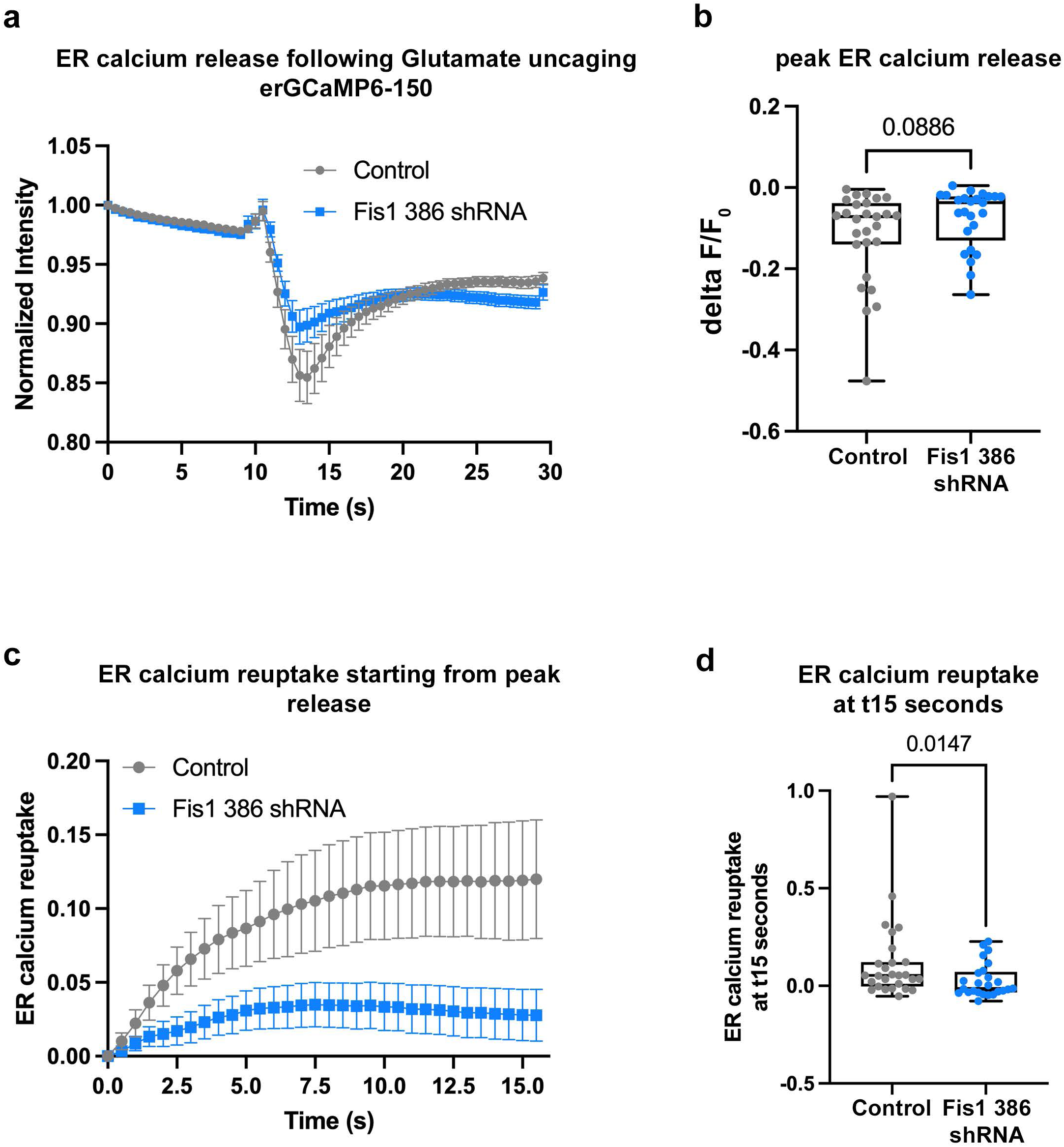
Fis1 knockdown reduces ER uptake following evoked neuronal activity. **a**) Plot of normalized erGCaMP6-150 intensity in control (grey) or Fis1 386 shRNA (blue) neurons following glutamate uncaging. **b**) Quantification of peak fluorescence decrease in control or Fis1 386 shRNA neurons showing slightly decreased, but not significant, ER calcium release in Fis1 KD neurons. **c**) Plot of normalized erGCaMP6-150 intensity in control (grey) or Fis1 386 shRNA (blue) neurons following peak release. **d**) Quantification of erGCaMP6-150 fluorescence recovery in control or Fis1 386 shRNA neurons showing decreased ER calcium re-uptake in Fis1 neurons. Control _erGCaMP6-150_ = 27 segments; Fis1 386 shRNA _erGCaMP6-150_ = 25 segments. p values are indicated in the figure following a Mann-Whitney test. Data are shown as individual points on box plots with 25^th^, 50^th^ and 75^th^ percentiles indicated with whiskers indicating min and max values.

**Supplementary Figure 5:**
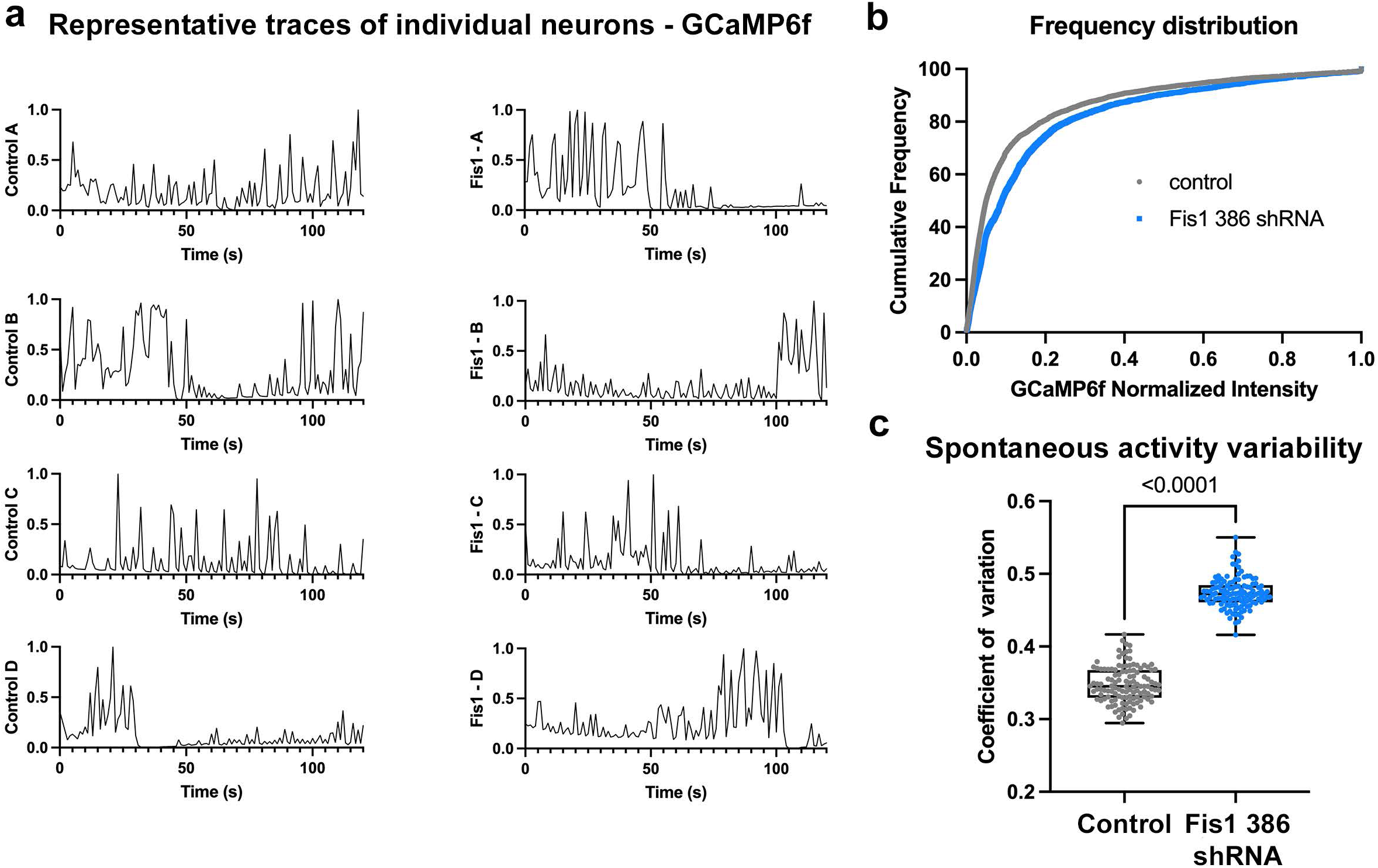
Fis1 knockdown doesn’t affect spontaneous neuronal activity. **a**) Representative traces of relative cytoplasmic GCaMP6f intensity to show spontaneous neuronal activity in 14DIV cultured cortical neurons electroporated with pCAG GCaMP6f and either control (left) or Fis1 386 shRNA (right). **b**) A cumulative frequency distribution of the normalized GCaMP6f intensities showing a rightward shift in Fis1 knockdown neurons. **c**) Quantification of the coefficient of variation demonstrating that Fis1 knockdown neurons have a higher variability in neuronal activity. Control _cyto GCaMP6f_ = 121 neurons; Fis1 386 shRNA _cyto GCaMP6f_ = 121 neurons. p value indicated in the figure following a Wilcoxon test. Data are shown as individual points on box plots with 25^th^, 50^th^ and 75^th^ percentiles indicated with whiskers indicating min and max values.

## Notes

### Competing Interest Statement

The authors have declared no competing interest.

